# Engineered production of isoprene from the model green microalga *Chlamydomonas reinhardtii*

**DOI:** 10.1101/2023.01.12.523746

**Authors:** Razan Z. Yahya, Gordon B. Wellman, Sebastian Overmans, Kyle J. Lauersen

**Affiliations:** Bioengineering Program, Biological and Environmental Sciences and Engineering Division, King Abdullah University of Science and Technology (KAUST), Thuwal 23955-6900, Kingdom of Saudi Arabia

**Keywords:** Isoprene, Microalgae, *Chlamydomonas reinhardtii*, MEP pathway, Volatile carbon

## Abstract

1.

Isoprene is a clear, colorless, volatile 5-carbon hydrocarbon that is one monomer of all cellular isoprenoids and a platform chemical with multiple applications in industry. Many plants have evolved isoprene synthases (IspSs) with the capacity to liberate isoprene from dimethylallyl pyrophosphate (DMAPP) as part of cellular protection mechanisms. Isoprene is hydrophobic and volatile, rapidly leaves plant tissues and is one of the main carbon emission sources from vegetation globally. The universality of isoprenoid metabolism allows volatile isoprene production from microbes expressing heterologous IspSs. Here, we compared heterologous overexpression from the nuclear genome and localization into the plastid of four plant terpene synthases (TPs) in the green microalga *Chlamydomonas reinhardtii*. Using sealed vial mixotrophic cultivation, direct quantification of isoprene production was achieved from the headspace of living cultures, with the highest isoprene production observed in algae expressing the *Ipomoea batatas* IspS. Perturbations of the downstream carotenoid pathway through keto carotenoid biosynthesis enhanced isoprene titers, which could be further enhanced by increasing flux towards DMAPP through heterologous co-expression of a yeast isopentenyl-PP delta isomerase. Multiplexed controlled-environment testing revealed that cultivation temperature, rather than illumination intensity, was the main factor affecting isoprene yield from the engineered alga. This is the first report of heterologous isoprene production from a eukaryotic alga and sets a foundation for further exploration of carbon conversion to this commodity chemical.

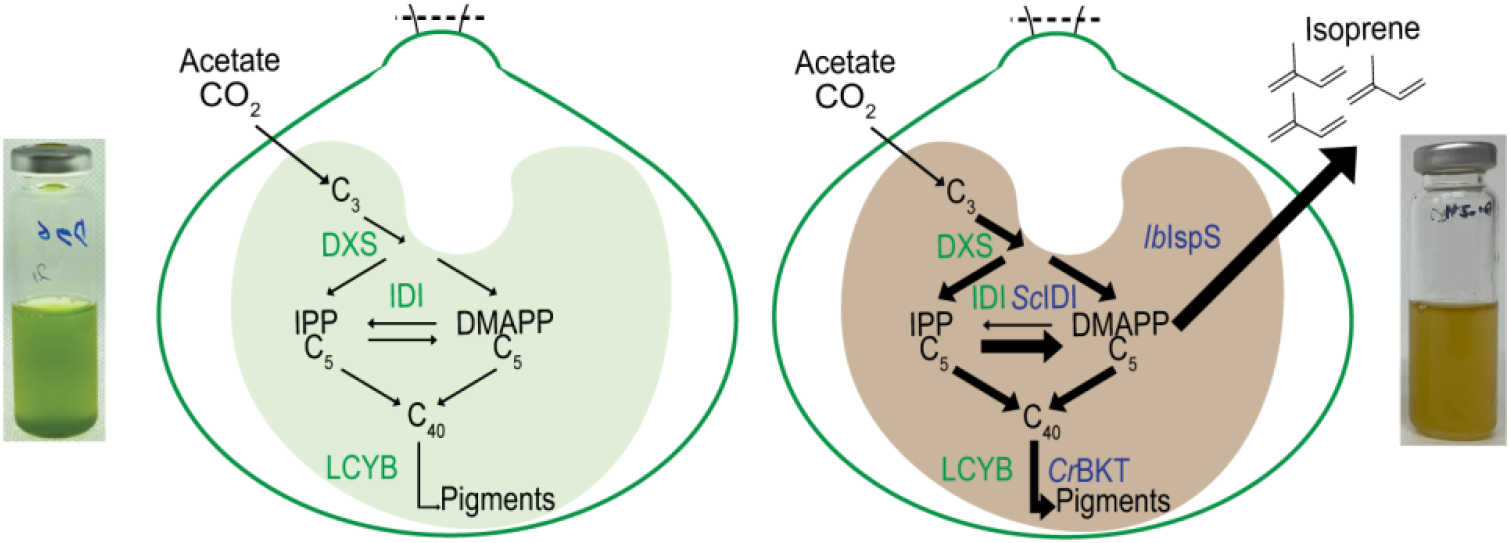

## 2. Introduction

Isoprenoid metabolism is required to generate carbon skeletons used to meet various needs of cellular processes and its presence can be found across all domains of life (Lange et al., 2000). Universal isoprenoid prenyl di-phosphates serve as substrates for chemical specialization in different organisms through the evolution of terpene synthases (TPs) and cytochrome P450s which together form complex chemical structures for various roles (Andersen-Ranberg et al., 2016; Pateraki et al., 2017, 2014). The universality of isoprenoid precursor availability across the domains of life has enabled demonstration that modular pathways producing specialized terpenoid chemicals can be transferred from a progenitor organism to another (Chandran et al., 2011). Heterologous isoprenoid production has now been shown in multicellular, unicellular, prokaryotic, and eukaryotic organisms (Andersen-Ranberg et al., 2016; Kirby and Keasling, 2009; Lauersen, 2019; Lindberg et al., 2010; López et al., 2020). Metabolic engineering in fermentative microbes is well established in bacteria and yeasts, organisms which consume organic carbon (sugar) and release carbon dioxide (CO_2_) through respiration. Fermentation via sugar consumption, requires the organic carbon substrate be produced by photosynthesis in a plant, refined, and shipped to site.

In contrast to their fermentative counterparts, phototrophic organisms are adapted to lifestyles of CO_2_ consumption as a source of carbon for growth (Melis, 2012). Powered by light energy, phototrophic organisms represent a greenhouse gas emission-less (carbon neutral) alternative to fermentative organisms and allow bio-production directly from CO_2_ conversion to biomass and target bio-products. To date, numerous metabolic engineering efforts have been reported in photosynthetic cyanobacteria (Chaves and Melis, 2018; Davies et al., 2015; Yunus et al., 2018; Yunus and Jones, 2018) while eukaryotic algae are also starting to take stage as phototrophic host organisms in which to demonstrate metabolic engineering concepts (Lauersen, 2019).

*Chlamydomonas reinhardtii* is a single-celled phototrophic model eukaryotic green microalga that can incorporate carbon into its biomass from either CO_2_ fixation through the Calvin-Benson-Bassham (CBB) cycle and/or acetic acid incorporation through the glyoxylate cycle (Lauersen et al., 2016b; Rochaix, 1995). *C. reinhardtii* and other green algae meet their entire isoprenoid needs solely from the 2-C-methyl-D-erythritol 4-phosphate/1-deoxy-D-xylulose 5-phosphate (MEP/DOXP) pathway localized in its plastid and has lost the mevalonate isoprenoid pathway through evolution (Lohr et al., 2012). The MEP pathway converts 3-carbon glycerol-3-phosphate (G3P) and pyruvate (Pyr) into the 5-carbon diphosphates dimethylallyl- and isopentenyl-pyrophosphate (DMAPP and IPP, respectively) (Schwender et al., 1996). These 5-carbon molecules are used to build higher-order carbon backbones that form cellular isoprenoids including sterols, carotenoids, and electron transfer components of enzymes, among many other roles (Kirby and Keasling, 2009; Lohr et al., 2012; Wichmann et al., 2020). Metabolic engineering in *C. reinhardtii* for the production of higher plant sesqui- (C15) and diterpenoids (C20) has been achieved by localization of TPSs in its cytoplasm and plastid, respectively (Einhaus et al., 2022; Lauersen et al., 2018, 2016a; Wichmann et al., 2018). Isoprene (C_5_H_8_, 2-methyl-1,3-butadiene) is a hemiterpene (C5) hydrocarbon that is produced by directly releasing the hydrocarbon backbone from the pyrophosphate groups of dimethylallyl pyrophosphate (DMAPP), a reaction catalyzed by isoprene synthases (IspSs) (Ilmén et al., 2015; Miller et al., 2001). Isoprene is clear and colorless in liquid form, volatile at 34 °C, and rapidly evaporates at room temperature or from biological tissues (Melis, 2012; Vickers et al., 2009a). Many, but not all, plants have evolved the capacity to produce isoprene to increase their tolerance towards reactive agents (Vickers et al., 2009b, 2009a). In some plants, as much as 20% of fixed carbon can be lost to volatile isoprene production and it is the major carbon emission product from vegetation globally (Sharkey et al., 2008; Sharkey and Loreto, 1993). Isoprene is also a commodity chemical used in the industrial formation of elastomers, rubber products, and chemical synthesis. Polyterpenoid rubbers (caoutchouc) can be sourced from the *Hevea brasiliensis* (rubber tree) and some species of dandelions (Pütter et al., 2017; Stolze et al., 2017); however, natural sources are not able to meet market demands. Currently, synthetic isoprene is sourced from several petroleum chemical refinement processes (Morais et al., 2015), but significant interest also lies in biological sources of isoprene production from microbial engineering (Matos et al., 2013).

Metabolic engineering of isoprene production in heterologous hosts has been investigated since the characterization of the first isoprene synthases (Chaves et al., 2017; Chaves and Melis, 2018; Miller et al., 2001; Vickers et al., 2017, 2014). The ability to transfer isoprene synthase genes into cell chassis allows the production of volatile isoprene from a microbial culture in scaled cultivation concepts and its harvest from the gaseous headspace phase above the fermentation broth (Whited et al., 2010). Reports have shown numerous isoprene synthases from different plants have different catalytic activities (Ilmén et al., 2015). These enzymes can be used in combination with tweaks to the MEP (Englund et al., 2018) pathway and/or with combinations of heterologous expression of the mevalonate pathway in organisms which do not naturally contain it (Bentley et al., 2014; Kim et al., 2016; Yang et al., 2012; Zurbriggen et al., 2012). Many insights have been gained into the mechanisms of carbon flow towards isoprenoids in plants as well as methods for improved titers and yields of molecular isoprene in microbial engineering concepts.

Here, we investigated whether *C. reinhardtii* could serve as a heterologous host for phototrophic isoprene production. We chose this eukaryotic alga due to availability of molecular tools and its ability to consume acetic acid mixotrophically, which allowed its cultivation in sealed headspace vials without metabolic waste toxicity. The expression of four plant IspSs from the algal nuclear genome and subcellular targeting to the plastid were investigated. It was determined that isoprene could be generated from the engineered alga and that the *Ipomoea batatas* IspS exhibited superior activity *in alga* over other IspSs. We then tested the effect of perturbations in the up- and downstream isoprenoid pathways on isoprene production. Modification of the terminal carotenoid pathway by the production of keto-carotenoids significantly improved isoprenoid production from the algal plastid, perhaps by increasing flux through the MEP pathway. It was also observed that an improved DMAPP-IPP ratio through heterologous expression of a yeast isopentenyl-diphosphate delta-isomerase (IDI) could complement the production increases from pigment modification to generate robust isoprene yields from the eukaryotic algal host. This is the first report of an engineered eukaryotic alga producing volatile isoprene and sets a platform for future bioprocess development wherein CO_2_ is converted into this platform chemical from photo-autotrophically grown green algae.

## 3. Materials and Methods

### 3.1. Algal culture conditions

*C. reinhardtii* strain UPN22 (Abdallah et al., 2022) was used for all experiments in this work. The alga was cultivated on solidified agar TAPhi-NO_3_ medium at 150 µmol m^-2^ s^-1^ light intensity or in 50 mL liquid TAPhi-NO_3_ medium shaken at 120 rpm (media recipes in (Abdallah et al., 2022)). UPN22 is a derivative of strain UVM4 (Neupert et al., 2009) that has been engineered to express the *Pseudomonas stutzeri* WM88 phosphite oxidoreductase (*ptxD*) (Loera-Quezada et al., 2016, p.) from its plastid genome (Changko et al., 2020) and its nitrate metabolism was complemented by transformation of native nuclear genes *nit1/nit2* (Fernández et al., 1989; Schnell and Lefebvre, 1993). UPN22 is able to grow with phosphite as the sole source of phosphorous and nitrate as a nitrogen source. These modifications can mitigate contamination and enable reliable growth of the alga in unfavorable conditions (Abdallah et al., 2022).

### 3.2. Genetic tool design, synthesis, transformation, and screening

Genes designed for expression in this work were adapted to the requirements of expression from the *C. reinhardtii* nuclear genome. Briefly, amino acid sequences were back-translated to most frequent codon usage for the nuclear genome of the alga and the first intron of *C. reinhardtii* ribulose bisphosphate carboxylase small subunit 2 (RBCS2i1) was systematically spread through each sequence *in silico* using the Intronserter program as previously described (Baier et al., 2020, 2018; Jaeger et al., 2019). Gene synthesis and subcloning were performed by Genscript (Piscataway, NJ, USA). The synthetic transgenes were cloned into pOpt3 expression plasmids (Gutiérrez et al., 2022) which contained the respective fluorescence reporters and selection markers indicated. For all genes, predicted plastid-targeting peptides were removed from the amino acid sequence prior to *C. reinhardtii* nuclear genome optimization. Four plant isoprene synthases (IspS) were genetically redesigned for algal expression in this work: *Populus alba* (UniProt Q50L36), *Pueraria montana* (Kudzu (UniProt: Q6EJ97)), *Eucalyptus globulus* (NCBI: BAF02831.1 (Englund et al., 2018; Gao et al., 2016)), and *Ipomoea batatas* (NCBI: AZW07551.1, (Ilmén et al., 2015)). Translated IspS peptide sequences were searched against *C. reinhardtii* genome v5.6 (JGI – Mycocosm (Nordberg et al., 2014). Search parameters: Database Blastp - *C. reinhardtii*_202001117 filtered model proteins – e-value: 1.0E-5), no native sequences with significant similarity were identified.

The *Saccharomyces cerevisiae* isopentenyl-diphosphate delta-isomerase (*Sc*IDI) and *Daucus carota* lycopene beta-cyclase (*Dc*LCYB, NCBI: NP_001316089.1) were also synthetically redesigned here while our previously reported *Salvia pomifera* 1-deoxy-D-xylulose 5-phosphate synthase (*Sp*DXS, NCBI: AXL65958.1 (Lauersen et al., 2018)) and *C. reinhardtii* beta-carotene ketolase (*Cr*BKT, UniProt: Q4VKB4.1 (Perozeni et al., 2020; Pivato et al., 2021)) were also used. Final annotated sequences of all expression plasmids are provided in Supplemental File 01.

Nuclear transformations of UPN22 with plasmid DNA were performed following the glass bead protocol as previously described (Kindle et al., 1989). For each plasmid, 10 µg of DNA was linearized with *Xba*I+*Kpn*I in 100 µl reactions prior to transformation, algal cells were left to recover in TAPhi+NO_3_ liquid medium for 6 h before plating on selective medium (spectinomycin 200 mg L^-1^, paromomycin 15 mg L^-1^, and/or bleomycin 10 mg L^-1^) and left under constant illumination for ∼7 days prior to colony picking. Up to 384 colonies per transformation event were transferred to TAPhi-NO_3_ agar plates using a PIXL robot (Singer Instruments, UK). After a further three days, colonies were replicated by stamping using a ROTOR robot (Singer Instruments, UK) to new plates as well as plates containing 200 mg L^-1^ amido-black for fluorescence screening as previously described

(Abdallah et al., 2022; Gutiérrez et al., 2022). The *Cr*BKT was directly fused to the spectinomycin resistance gene *aadA* as previously reported (Pivato et al., 2021) and was screened by the color change of cells from green to brown due to the accumulation of ketocarotenoids as previously described (Perozeni et al., 2020). Other target transgenes of interest were expressed as a fusion to either mScarlet (red fluorescent protein (RFP), (Bindels et al., 2017)) or mVenus (yellow (YFP), (Kremers et al., 2006)). Screening of transformants for transgene expression was performed by fluorescence imaging at the agar-plate level to semi-quantitatively identify robust gene-of-interest-FP fusion protein expression as previously described (Gutiérrez et al., 2022). Briefly, chlorophyll fluorescence was captured to normalize colony presence/absence on plates with 475/20 nm excitation and DNA gel emission filter (640/160 nm) for 1 s exposure. Red fluorescence from RFP was captured with 560/10 nm excitation and 600/10 nm emission filters for 2:30, while YFP signals were captured with 504/10 nm excitation and 530/10 nm emission filters for 30 s (Analytik Jena Chemstudio Plus gel doc). Transformants with robust FP signals were selected and inoculated in 12-well plates in 2 mL of liquid TAPhi-NO_3_ medium, shaking at 160 rpm. Expressed protein-predicted molecular masses were then confirmed by sodium dodecyl sulphate-polyacrylamide gel electrophoresis (SDS-PAGE) in-gel fluorescence against the fusion protein fluorescent reporter as previously described (Gutiérrez et al., 2022).

### 3.3. Characterization of isoprene production in transformants

Colonies were taken from agar plates and incubated in 1 mL liquid TAPhi-NO_3_ medium in 24-well microtiter plates for 3 days, with constant light and 190 rpm orbital shaking. Then, replicates were transferred into 12-well plates containing 2 mL TAPhi-NO_3_ per well, and shaken at 160 rpm until the populations reached late logarithmic growth phase. 1 mL of each dense culture was added to 9 mL of the same medium in 20 mL autoclaved headspace bevel top vials with PTFE/silicon septum lids (Supelco, Cat.: 27306) which were then sealed. The culture-containing vials were kept standing and left in a 12 h:12 h light:dark cycle at 100 μmol photons m^−2^ s^−1^ for 6 d. To check whether the presence of isoprene in the headspace of a sealed vial affected the culture growth. A toxicity test was performed by adding 9 mL of media containing *C. reinhardtii* and 1 mL of known isoprene concentrations followed by 6 d cultivation in same condition as above.

For light and temperature trials, a total of 80 sealed GC vials containing algal culture were prepared as above and subsequently transferred into 20x 1L-Erlenmeyer flasks (n= 4 vials per flask) each filled with 800 mL of milliQ-H₂O with a PH probe to record the temperature. The flasks containing floating vials were kept in 20 separate Algem photobioreactors (Algenuity, UK) without shaking for 6 d before headspace sampling. The reactors were programmed to four different light intensities (70, 130, 190, 250 μmol photons m^−2^ s^−1^ as measured at culture distance from LEDs) and five temperatures (18, 21, 24, 27, 30 °C), thereby generating 20 distinct temperature-light combinations. The light spectra of the cultivation conditions are shown in Supplemental File 02.

Cell density measurements for all sealed cultivations were conducted by inverting the GC vials and sampling 0.5 mL of liquid culture from each vial through the septum using a 3 mL syringe and needle. Cell densities were analyzed by flow cytometry using an Invitrogen Attune NxT flow cytometer (Thermo Fisher Scientific, UK) equipped with a 488 nm blue excitation laser, a 488/10 nm emission filter for forward-scatter (FSC), and a 695/40 nm filter (BL-3) for chlorophyll fluorescence. Prior to analysis, each biological sample was diluted 1/100 with 0.9% NaCl solution and measured in technical triplicates following a previously described method (Overmans and Lauersen, 2022).

### 3.4. Headspace Gas Chromatography

Isoprene production inside sealed vials was quantified using headspace gas chromatography-mass spectrometry and flame ionization detection (HS-GC-MS-FID). The system used was an Agilent 7890A gas chromatograph connected to a 5977A inert MSD with triple-axis detector, equipped with a DB-624 column (60 m × 0.25 mm i.d., 1.40 μm film thickness) (Agilent J&W, USA). Prior to sample injection, the vials containing algal culture were incubated in an HT3 Static and Dynamic Headspace autosampler (Teledyne Tekmar, USA), which offers both static and dynamic analysis.

HS-GC-MS-FID analyses of samples from engineered algal strains were conducted using the static autosampler option. The vials were incubated at 40 °C for 20 min before a 2 min vial pressurization, 0.2 min pressurization equilibrium at 10 PSIG, 2 min loop fill at 5 PSIG, and 1 min sample injection. Headspace samples from parental strains were analysed using the dynamic autosampler mode. The vials were preheated for 15 mins at 40°C, followed by 5 min sweep flow at 50 mL min^-1^, 1 min dry purge at 25°C at 50 mL min^-1^, and trap bake was done for 5 min at 230°C at a rate of 150 mL min^-1^. For all samples, the initial GC oven temperature was 50 °C, then raised to 180 °C at a rate of 10 °C min^−1^, followed by 25 °C min^−1^ to 200 °C, resulting in a total GC cycle time of 15 min at a constant flow of 1mL/min with helium as carrier gas. Gas chromatograms were evaluated with MassHunter Workstation software v. B.08.00 (Agilent Technologies, USA). The National Institute of Standards and Technology (NIST) library (Gaithersburg, MD, USA) was used to identify isoprene, ethanol, argon, and CO_2_. Immediately before each GC run, serial dilutions of 99% pure Isoprene (Sigma-Aldrich) in TAPhi-NO_3_ medium were prepared in triplicate (n=3) at five different concentrations for the engineered strains (10–500 ppm) and parental strains (0.5–3 ppm). Standards were analyzed in the same way as samples, and isoprene peak areas were plotted against standard concentrations to generate a standard curve for isoprene quantification.

### 3.5. Data analysis

To determine whether mean volumetric isoprene production (mg L^-1^ culture) and cell-specific isoprene production (pg cell^-1^) varied significantly between *C. reinhardtii* cultivated in the 12 h:12h light:dark cycle and in 24 h light, we performed a total of six Student’s *t*-tests (one test per parameter per plasmid). Similarly, to identify if mean isoprene production was significantly different between parental strains and engineered strains, a total of 28 individual Student’s *t*-tests (14 tests for volumetric isoprene production and 14 tests for cell-specific production) were carried out. All statistical analyses were performed using the software JMP Pro 16.2 (SAS Institute Inc., Cary, NC). Means were considered significantly different at a level of *p* < 0.05.

## 4. Results and Discussion

### 4.1. Chlamydomonas as a host for heterologous isoprenoid production

As alternatives to fermentative hosts, algal chassis can produce heterologous products of interest directly from CO_2_ as a carbon source, while consuming nitrogen and phosphorous sources in water (de Freitas et al., 2023). These properties make microalgae interesting for sustainable bio-production concepts that convert waste to value. Post-treatment waste waters contain nitrogen and phosphorous that should be removed prior to emission to the environment or reuse, and CO_2_ is a common waste emission from industrial and human processes. Compared to other model organisms, *C. reinhardtii* has been delayed in its use for demonstration of complex metabolic engineering due to a known recalcitrance of transgene expression from its nuclear genome (Lauersen, 2019). A silencing mechanism acts upon transgenes as they integrate into the nuclear genome, which has previously hindered efforts to overexpress heterologous transgenes in this host (Neupert et al., 2020). Expression of transgenes has also been found to depend on complex interactions of intron sequences and host regulation machinery, in addition to high-GC content and specific codon bias requirements (Baier et al., 2018; Eichler-Stahlberg et al., 2009; Fuhrmann et al., 1999; Lumbreras et al., 1998). We previously demonstrated that endogenous introns spread in specific patterns throughout synthetically optimized transgenes enables their robust expression from the nuclear genome of this alga and that having exon regions longer than ∼500 bp results in poor transgene expression (Baier et al., 2020, 2018). The development of strains with mutations in the Sir2-type histone deacetylase (SRTA) (Neupert et al., 2020, 2009) coupled with strategic transgene designs that work with the host regulation machinery (Jaeger et al., 2019) has enabled the awakening of *C. reinhardtii* as a versatile chassis for metabolic engineering studies (Schroda, 2019).

Many reports now exist demonstrating the capacity of heterologous isoprenoid production in *C. reinhardtii* (Einhaus et al., 2022; Lauersen et al., 2018). It has been suggested, that as a photosynthetic organism, flux through its isoprenoid pathways should be highly adaptable to the pull of heterologous isoprenoid production due to the needs of replenishing pigments involved in light-harvesting. In support of this notion, heterologous production of sesqui- and diterpenoids have been shown without impacting native pigment profiles of the alga (Einhaus et al., 2022; Lauersen et al., 2018). Through re-design and overexpression of its native beta-carotene ketolase (*Cr*BKT), it was also shown that keto-carotenoids can be produced in vegetative cells of *C. reinhardtii*, their intense red-orange colors producing a distinctive brown color in combination with algal chlorophylls in living cells (Cazzaniga et al., 2022; Perozeni et al., 2020). Extreme perturbation of the carotenoid profile could be seen as a metabolically expensive phenotype for the algal cell; however, it has been shown that *Cr*BKT overexpression lines exhibit improved high-light tolerance and growth (Cazzaniga et al., 2022). Although the alga relies entirely on the MEP pathway for isoprenoid generation, it likely has a distinct capacity to adapt to metabolic pulls of further metabolic engineering for heterologous isoprenoid production. In this context, we were interested in the ability of *C. reinhardtii* to produce volatile isoprene as a co-product to its biomass.

### 4.2. Chlamydomonas can be engineered to produce high titers of isoprene

*Chlamydomonas* can grow photo-litho-autotrophically in light with CO_2_, photo-heterotrophically and heterotrophically with acetic acid, and in combination with acetic acid and CO_2_ as carbon sources. We took advantage of this behavior to grow cells in sealed vials, using acetic acid as a carbon source and light for energy (Fig. 1A). The parental strain was first tested for growth in sealed vials and, consequently, all further isoprene production tests were performed in the same way (Fig. 1A). In this condition, the cells obtain most of their energy from light, consume acetic acid as a carbon source, and metabolic CO_2_ produced by respiration is fixed by the CBB cycle. CO_2_ was not detected in GC-MS-FID analysis in any condition of acetic acid growth of the alga alone (Supplemental Fig. 1). UPN22 cells grew in sealed vials with a 50:50 ratio of culture medium to headspace (Fig. 1A). By the 7^th^ day of cultivation, only minute traces of isoprene were detectible from the parental UPN22 strain (∼40 mg L^-1^ (0.04 ppm), Fig. 1B). The addition of pure isoprene to cultures resulted in a non-linear effect on cell accumulation but did not abolish growth in completely sealed vials; even at concentrations of up to 1 g L^-1^ (Fig. 1C).

**Figure 1.**
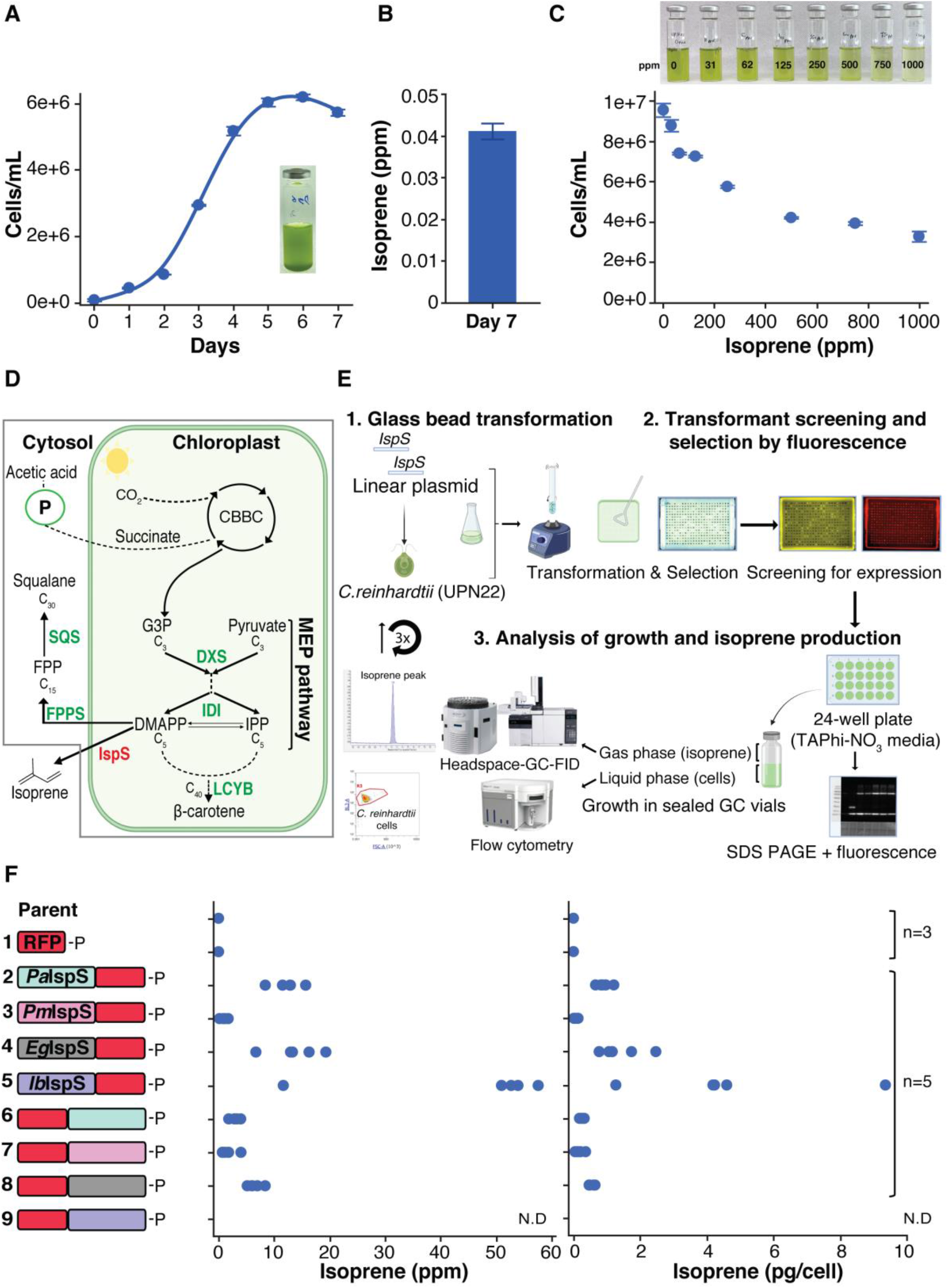
*C. reinhardtii* UPN22 parental (A) cell concentration over time and (B) isoprene production on day 7 when grown photo-mixotrophically in sealed GC-vials (pictured). (C) Isoprene toxicity panel showing a decline in cell density with increasing isoprene concentration added to sealed culture vials. (D) Pathways involved in isoprene metabolism in the green algal cell. (E) Laboratory workflow used as part of this study. (F) Volumetric and per-cell production of isoprene in the parent strain and nine other groups of transformants generated with different IspS expression plasmids.

The alga maintains the MEP pathway in its plastid where it supplies precursor prenylphosphates for carotenoid pigments and, in the cytosol, the C15 precursors of sterol biosynthesis (Fig. 1D) (Lohr et al., 2012; Wichmann et al., 2020). Carbon is supplied to the MEP pathway as glycerol-3-phosphate (G3P) and pyruvate (Pyr), which come from carbon dioxide fixed in the CBB cycle or acetic acid uptake through the glyoxylate cycle partially via the peroxisomes (Fig. 1D) (Lauersen et al., 2016b). Heterologous expression of isoprene synthases, targeted to the algal plastid, should use inherent DMAPP as substrate to yield molecular isoprene (Fig. 1D). To engineer expression of heterologous IspSs in *C. reinhardtii*, genes for four plants: *Populus alba* (white poplar), *Pueraria montana* (Kudzu), *Eucalyptus globulus* (southern blue gum), and *Ipomoea batatas* (sweet potato), were synthetically optimized and cloned into expression plasmids in fusion with the mScarlet (red fluorescent protein (RFP)) in both N- and C-terminal fusion orientations (described in Methods, sequences in Supplemental File 1). These plasmids were transformed into the alga and fluorescence screening was used to identify fusion protein expression in isolated transformants (Fig. 1E). Correct molecular mass of each fusion protein was observed by in-gel SDS-PAGE fluorescence analysis against the RFP (Fig. 1E, Supplemental Fig. 2). For each plasmid, 5 confirmed transformants were subjected to growth in sealed GC vials, followed by analyses of isoprene accumulation and cell density measurements on cultivation day 6 by headspace GC-MS-FID and flow cytometry, respectively. Volatile isoprene was readily detected in culture vials of all four IspSs in the ppm range (mg L^-1^) after 6 d sealed cultivation (Fig. 1F). We chose day 6 for these analyses as 8-day growth tests with two *Ib*IspS-RFP expressing transformants showed cell density and isoprene accumulation reached its maximum here and cell counts were unreliable after this day (Supplemental Fig. 3). The orientation of fusion protein greatly affected isoprene production, positioning at the C-terminus of the RFP resulted in lower titers for all constructs. No transformants could be obtained from RFP-*Ib*IspS plasmids (Fig. 1F, N.D.). Volumetric yields of isoprene followed cellular productivities with production rates from highest to lowest across constructs with *Ib*IspS > *Eg*IspS > *Pa*IspS > *Pm*IspS (Fig. 1F). Recent reports have shown that both *Eg*- and *Ib-*IspSs are catalytically superior to those previously reported (Englund et al., 2018; Gao et al., 2016; Ilmén et al., 2015). *Eg*IspS was used to demonstrate gram-scale production of isoprene from *Synechococcus elongatus* when it was fused directly to an IDI (Gao et al., 2016). Both enzymes required modified precursor flux through IDI or DXS overexpression before being stable in *Synechocystis* sp. PCC 6803 (Englund et al., 2018). In its initial report, *Ib*IspS produced more isoprene than all other tested IspSs in *Escherichia coli* (Ilmén et al., 2015; Li et al., 2019). In *C. reinhardtii, Eg*IspS was by far the lowest expressed of the four IspSs, with lower fluorescence signals during screening than the other three plant enzymes (not shown). Despite its lower expression, *Eg*IspS produced more isoprene than the *Pa*IspS, which has long been used as an efficient synthase in fermentative production of heterologous isoprene (Vickers et al., 2009b; Whited et al., 2010; Yang et al., 2012). For *Ib*IspS, 4 out of 5 transformants investigated produced between 50–58 ppm (mg L^-1^) isoprene in their sealed vial in 6 days cultivation in TAPhi-NO_3_ medium (Fig. 1F). Therefore, we looked to further investigate whether isoprene production could be improved in *C. reinhardtii* by perturbing up- and downstream reactions in the MEP and carotenoid pathways of the algal host expressing this synthase.

### 4.3. Pigment perturbation & DMAPP:IPP ratio affects algal isoprene production

The precursor of isoprene, DMAPP, is one of two C5 isomers (IPP and DMAPP) produced by the MEP pathway on route to higher-carbon atom containing isoprenoids (Lohr et al., 2012; Schwender et al., 1996). For longer chain isoprenoid production, more IPP is needed than DMAPP, which may bias evolution of metabolic ratios of these two isomers in different organisms. DMAPP and IPP ratios were previously thought to be balanced at a 1:5 ratio in all organisms. However, it is now known that this ratio is organism-specific and related to activity of the terminal enzyme of the MEP pathway hydroxymethylbutenyl diphosphate reductase (HDR) (Bongers et al., 2020). Prenyltransferases: geranyl-, farnesyl-, and geranylgeranyl pyrophosphate synthases (GPPS, FPPS, and GGPPS, respectively) build higher chain length isoprenoids from IPP and DMAPP and have much faster catalytic rates than known IspSs (see: brenda-enzymes.org). DMAPP, therefore, sits as a building block in the middle of the isoprenoid metabolic pathway in a photosynthetic cell, produced from upstream reactions of the MEP pathway and pulled from downstream reactions of the sterol and chlorophyll/carotenoid pathways.

To produce isoprene, an IspS must compete with the pull of competitive enzymes, while also requiring that the supply of substrate match its catalytic activity. To determine if isoprene titers from *C. reinhardtii* could be increased, here, both the up- and down-stream isoprenoid pathways were perturbed in a strain expressing the *Ib*IspS. Increasing *Ib*IspS titer by additional transformation and integration events was also investigated as this strategy has been shown important to increase TPS titers for sesqui- and diterpene synthases in the algal cytoplasm and plastid (Einhaus et al., 2022; Lauersen, 2019; Wichmann et al., 2022, 2018).

Up-stream perturbations investigated here employed expression of a de-regulated 1-deoxy-D-xylulose 5-phosphate synthase (DXS) variant from *S. pomifera* (*Sp*DXS), the first rate-limiting enzyme of the MEP pathway. In addition, the ratio of DMAPP and IPP was modified to bias DMAPP accumulation through heterologous expression of the yeast isopentenyl-diphosphate delta-isomerase (*Sc*IDI). Downstream carotenoid biosynthesis was perturbed by heterologous overexpression of the carrot lycopene beta-cyclase (*Dc*LCYB) or overexpression of the native *C. reinhardtii* beta-carotene ketolase (*Cr*BKT). The expression of these four heterologous genes and localization to the algal plastid did not result in isoprene formation without an IspS present (Fig. 2A, B). The relative positions of each enzyme in the native MEP and carotenoid pathways are shown in Fig. 2C.

**Figure 2.**
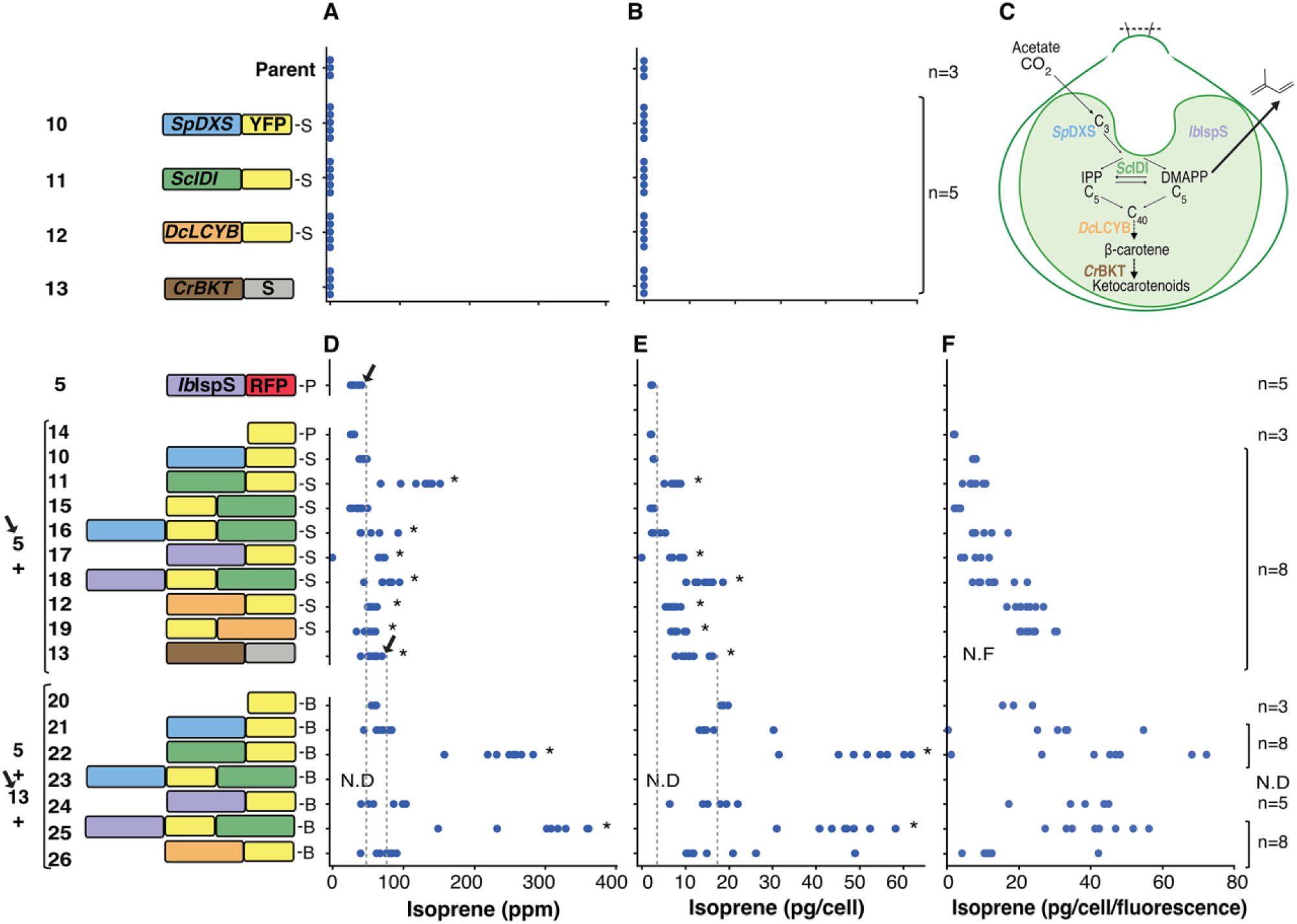
Volumetric- (A) and per cell- (B) isoprene production for *C. reinhardtii* UPN22 parental strain and transformants with heterologous genes used to perturb the MEP and carotenoid pathways without *Ib*IspS. localization of each heterologous gene expressed and the native MEP pathway. Volumetric- (D), per cell- (E), and per-cell per-fluorescence (F) isoprene production for *Ib*IspS transformant (5), transformed with ten spectinomycin resistance (-S) plasmids and *Ib*IspS+*Cr*BKT dual transformant (5+13) transformed with seven bleomycin resistant (-B) plasmids expressing different target enzymes. N.D. indicates isoprene not detected; N.F. indicated no fluorescence detected; n indicates number of unique transformants analyzed; Asterisks in D and E indicate instances where mean volumetric isoprene titers and cell-specific isoprene production, respectively, were significantly different from the means of the YFP control (Student’s *t*-tests, *p* < 0.05).

An *Ib*IspS-RFP expressing transformant (Fig. 2D, plasmid 5, black arrow) was subsequently transformed with a series of plasmids expressing each new gene of interest fused to YFP and conferring resistance to spectinomycin (Fig. 2D, E, F). Transformants expressing plasmid 5 in addition to the second plasmid were selected on paromomycin and spectinomycin containing medium as the original *Ib*IspS plasmid conferred paromomycin resistance. Transformation and expression of plastid targeted YFP alone did not improve isoprene yields in the *Ib*IspS-RFP expressing line, as anticipated (Fig. 2D, E, F - plasmids 5+14). Expression of *Sp*DXS did not improve titers of isoprene volumetrically or per cell, while N-terminal *Sc*IDI-YFP was found to significantly improve isoprene titers to 68–152 mg L^-1^, or 5–9 pg cell^-1^ (Fig. 1D, E, respectively - plasmids 5+11). The combination of *Sp*DXS-YFP-*Sc*IDI in one plasmid had significantly improved volumetric titers, but not when normalized to cellular productivity (Fig. 2 D, E - plasmids 5+16). Significant isoprene titer increases were also observed from *Ib*IspS-YFP or *Ib*IspS-YFP-*Sc*IDI fusion proteins, with the later strains exhibiting isoprene yields between 7–19 pg cell^-1^ (Fig. 2D, E, F - plasmids 5+17,18).

Perturbations of the carotenoid pathway at the level of lycopene beta-cyclase (LCYB) have been previously shown to increase overall carotenoid titers in several plants, an effect attributed to an overabundance of apocarotenoid hormones and consequent growth regulation (Moreno et al., 2020). Overexpression of a phytoene-β-carotene synthase from yeast with LCYB activity in *C. reinhardtii* has also been reported to improve beta carotene titers in algae (Rathod et al., 2020), however, it was unclear if heterologous perturbation of carotenoid pools would effect isoprene titers produced from an *Ib*IspS expressing strain. Both fusion orientations of *Dc*LCYB to YFP (plasmids 12/19) were found to increase isoprene titers significantly to 6–10 pg cell^-1^ (Fig. 2D, E - plasmids 5+12,19). However, the most significant effect of carotenoid pathway modulation was seen when *Cr*BKT was co-expressed in the *Ib*IspS-RFP transformant which resulted in isoprene titers of 40–70 mg L^-1^ and 8–16 pg cell^-1^ (Fig. 2D, E - plasmids 5+13). Cells transformed with this plasmid were screened for the presence of ketocarotenoids by the brown color change previously described (Perozeni et al., 2020) as well as the maintenance of the RFP signal of the *Ib*IspS-RFP (plasmid 5).

Plasmid 13 does not contain a YFP reporter as the *Cr*BKT is directly linked as a fusion protein to the spectinomycin resistance gene (*aadA*, Supplemental File 1). The antibiotic resistance gene (*aadA*) of the YFP containing gene expression cassettes of the above plasmids (10-19) was then replaced with the bleomycin/zeocin resistance gene (*Sh*Ble) using the modularity in the pOpt3 plasmid backbone. Newly generated plasmids 20–26 were then transformed into a *Ib*IspS-RFP + *Cr*BKT-*aadA* (plasmids 5+13) expressing transformant followed by selection on all three antibiotics. Colonies isolated with brown ketocarotenoid phenotype as well as red and yellow fluorescence from FP fusion proteins were again subjected to 6-days growth in sealed GC vials and checked for isoprene production in their headspace. Transformants from two plasmids stood out as exhibiting significant further improvements to isoprene yields compared to their progenitor strain (Fig. 2D, lower black arrows - plasmids 5+13). Transformants with plasmids 5+13 and *Sc*IDI-YFP were observed to produce 152– 283 mg L^−1^, 32–62 pg cell^−1^ while those transformed with *Ib*IspS-YFP-*Sc*IDI produced from 149–362 mg L^-1^ and 31–58 pg cell^-1^ (Fig. 2 D, E - plasmids 5+13+22 and 5+13+25, respectively).

The perturbations to the algal isoprenoid pathway tested here affected both up and downstream portions of its isoprenoid metabolism. Fig. 3 depicts proposed effects that each significant modification had on the algal metabolism and its engineered production of isoprene liberated from DMAPP. The parental strain converts CO_2_ or acetic acid into the C5 precursors IPP and DMAPP, where native *Cr*IDI should balance their pools (Fig. 3A). Expression of the heterologous *Ib*IspS created modest amounts of isoprene from available DMAPP, which is a small remaining pool of what is not rapidly converted by the cell into higher-order isoprenoids like pigments (Fig. 3B). Expression of the *Cr*BKT extends the carotenoid pathway to produce ketocarotenoids like canthaxanthin and astaxanthin (Fig. 3C). This likely perturbs the terminal regulation of the MEP pathway and causes an increased flux which results in a greater abundance of DMAPP as an intermediate for the active *Ib*IspS and results in improved isoprene titers (Fig. 3C). The *Ib*IspS appears to have a high catalytic activity compared to other IspSs. In a previous report, when Western blot signals were fainter for this enzyme expressed in a cyanobacterium, it produced similar titers to the *Eg*IspS which is considered a highly catalytically active IspS (Englund et al., 2018). The efficiency of *Ib*IspS was shown to be capable of producing even more isoprene when *Sc*IDI was added to strains with *Cr*BKT overexpression (Fig. 3D). The combination of simply *Sc*IDI alone (5+13+22) was comparable to a fusion of *Ib*IspS-YFP-*Sc*IDI, which also has increased abundance of the IspS. This suggests that DMAPP substrate availability is the limiting factor at this stage of engineering isoprene production from the alga. Engineering the cell to supply more DMAPP to the *Ib*IspS through both BKT mediated pull on the carotenoid pathway coupled to isomerase conversion of IPP to DMAPP is a combinatorial strategy to meet the catalytic potential of the *Ib*IspS.

**Figure 3.**
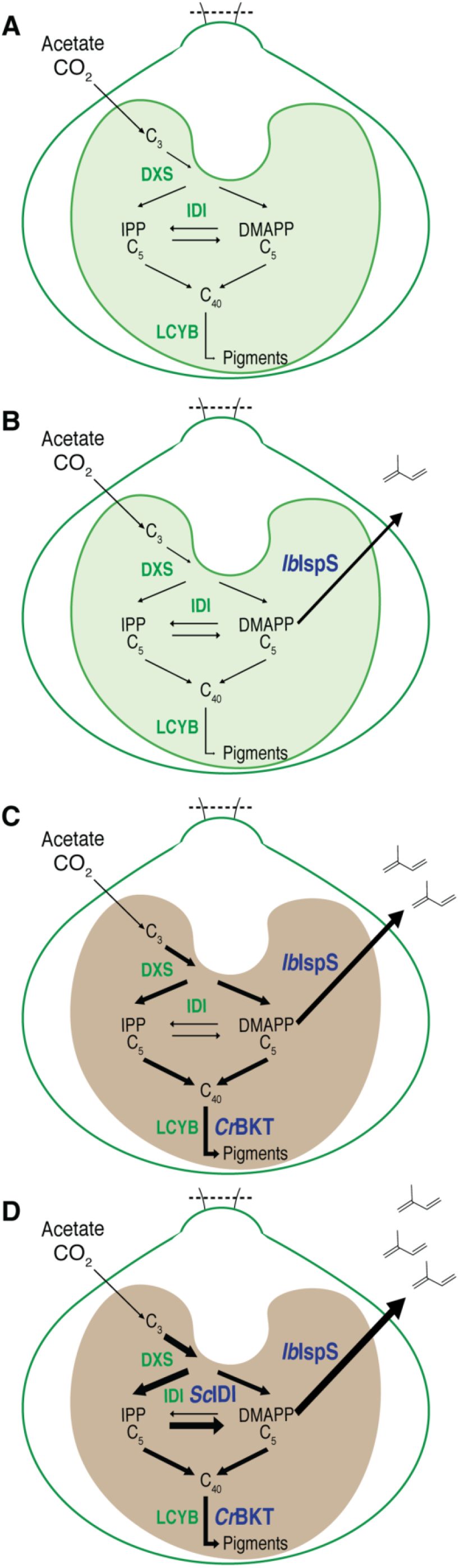
Modification of *C. reinhardtii* metabolism and its engineered production of isoprene- (A) parental strain. (B) parental strain with the expression of *Ib*IspSs. parental strain with the expression of *Ib*IspS and *Cr*BKT. (D) parental strain with the expression of *Ib*IspS, *Cr*BKT and *Sc*IDI.

### 4.4. Temperature affects engineered algal isoprene production more than illumination

Previous reports of sesqui- and diterpenoid production from *C reinhardtii* have shown that the production of isoprenoids can be improved if cell-growth is kept at modest levels (Einhaus et al., 2022; Lauersen et al., 2018, 2016a; Wichmann et al., 2022, 2018). This is seemingly improved for diterpenoids if day:night cycles are used rather than 24 hour light cycles (Einhaus et al., 2022; Lauersen et al., 2018). It is assumed that in the dark, heterologous terpene synthases have improved access to the prenylphosphate precursors, which would otherwise be strongly pulled for photosynthetic roles. The presence of ketocarotenoids in *C. reinhardtii* has been shown to improve aspects of light tolerance for the alga but also affect pigment ratios (Cazzaniga et al., 2022). Accumulation of ketocarotenoids in *C. reinhardtii* has been shown to reduce total chlorophylls and carotenoids, reorganize photosystems, and cause overall changes in the total carotenoid profile of the green alga (Cazzaniga et al., 2022; Perozeni et al., 2020). Therefore, it was important to determine if light regime affected isoprene production in the same manner across representatives of the different plasmid combinations. Transformant strains expressing Plasmid #5 alone have no ketocarotenoids, those with 5+13 have ketocarotenoid accumulation, and those with plasmids 5+12+25 have ketocarotenoids and modified DMAPP:IPP ratios (Figure 4). It was observed that for all combinations of plasmids, significantly higher cellular isoprene titers were produced in 24 h illumination cycles (Fig. 4A, B, C). This is in contrast to other classes of terpene synthases which seemingly produce more terpenoid products when dark cycles are employed.

**Figure 4.**
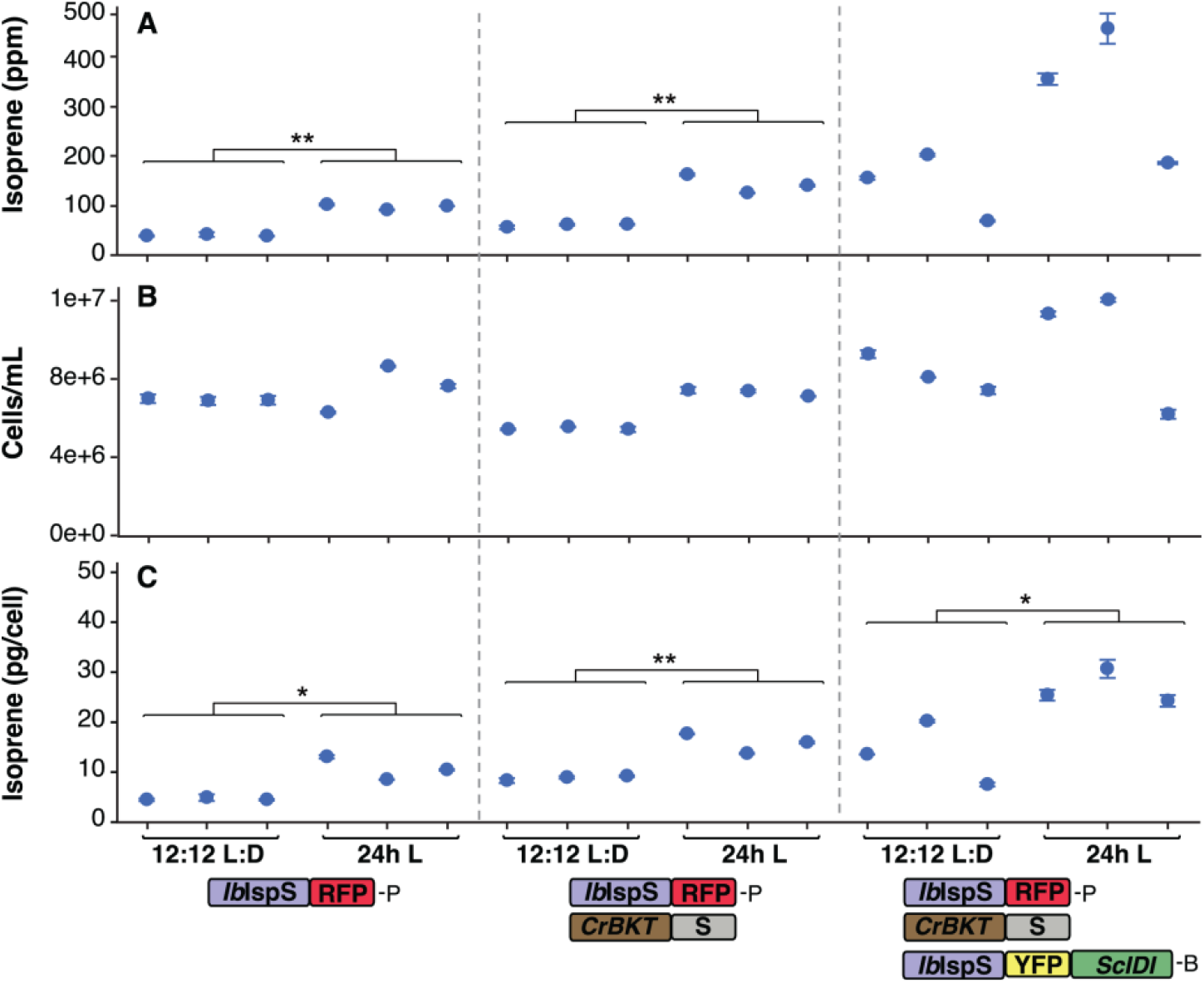
Three transformants per genetic construct were cultivated in 12:12 h day:night or 24-hour light cycles in biological triplicates to determine the impact of light cycle on isoprene productivity. Mean ± SEM (A) volumetric isoprene production, (B) *C. reinhardtii* cell concentration and (C) isoprene production per-cell for different plasmids and light cycles. Asterisks in panels A and C indicate the significance results of individual Student’s *t*-tests (* *p* < 0.05; ** *p* < 0.01).

It was recently shown that heterologous patchoulol production was lower when cells were exposed to modeled higher temperature and light environmental conditions (de Freitas et al., 2023), suggesting that both temperature and light regimes impact isoprenoid biosynthesis in the alga. Isoprene synthases have optimal catalytic activities at ∼42 °C (Li et al., 2019). *C. reinhardtii*, however, exhibits optimal growth between 28–30 °C (Stern et al., 2008). Temperature in culture can be directly affected by light irradiance, and it is difficult to separate the effect of these two factors on an algal culture. Therefore, we employed 20-parallel Algem photobioreactors to precisely control temperature and light levels and tease apart which of the two parameters has a greater effect on the engineered isoprene productivity from the alga. Using sealed headspace culture vials suspended in water inside photobioreactors, temperature and light could be independently controlled, and the temperature effect of irradiance mitigated by active cooling (Figure 5). Cultivations were conducted in an array of 20 combinations of temperature (18, 21, 24, 27, and 30 °C) and light conditions (70, 130, 190, 250 µE) to analyze behaviour across these two factors (Figure 5). We observed colorimetric changes in the phenotype of the ketocarotenoid producing strain, with lower light intensities resulting in more green-phenotypes while higher light exhibited brown-orange phenotypes (Fig. 5 A). Chlorophyll a, b and total carotenoids were highest in cells at the lowest light and temperature conditions and lowest at higher irradiances (Figure 5A). This is in line with recent reports of lower total pigments in higher light conditions for *Cr*BKT expressing lines of *C. reinhardtii* (Cazzaniga et al., 2022). Isoprene production followed a pattern across light intensities, that cultures grown at 30 °C exhibited the highest productivity regardless of light intensity (Figure 5B). Surprisingly, highest isoprene volumetric titers were observed in lower light intensities in the highest temperature conditions (70 µE, 27 and 30 °C, Figure 5B). This could suggest that despite lower total carotenoids per cell, the natural metabolic pull on IPP and DMAPP towards pigments during high-light is stronger than the catalytic capacity of *Ib*IspS to produce isoprene. It may also suggest that reduced carotenoids per cell in higher light also downregulates the flux through the MEP pathway to the intermediates and consequently makes less DMAPP available.

**Figure 5.**
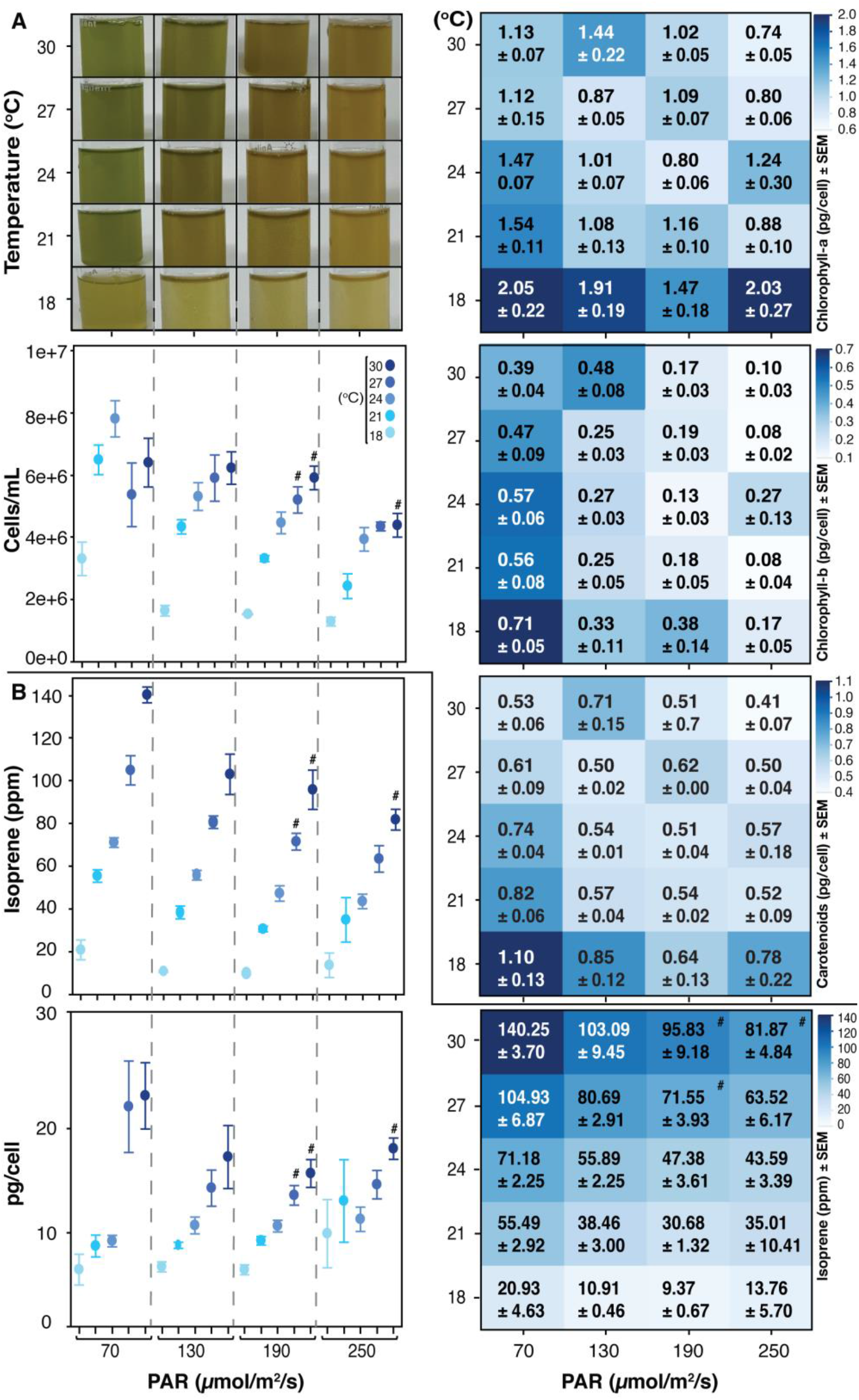
(A) Algal culture color, cell count, and pigment concentrations (chlorophyll-a, -b, and carotenoids) after 6 d cultivation in different temperature light combinations. (B) Volumetric (ppm – mg L^-1^) and per-cell (pg cell^-1^) isoprene production after 6 d. All values indicate means ± SEM (n=4 biological replicates) expect for samples labeled with hashtag (n=3).

For the green alga, catalytic activity of the *Ib*IspS mediated by temperature appears to be the most significant factor in isoprene production, and that optimal enzyme activity is likely outside its range of growth temperatures. However, daily high temperature peaks are tolerated for *C. reinhardtii*, as recently shown in environmental modelling in the extreme environment of the Red Sea coast (de Freitas et al., 2022). It could, therefore, be possible to create cultivation parameters that balance cell growth densities and temperature in lower and consistent light conditions to maximize isoprene yields. It must, however, be noted that these findings are from mixotrophically grown cultures, and the dynamics of the MEP pathway intermediates may be different when grown exclusively on CO_2_ as a carbon source. It has been previously shown that the carbon source directly affects per-cell chloroplast produced diterpenoid production in C. reinhardtii, unfortunately, it was not possible to supply CO_2_ to the sealed vial cultivations used in this test. Future efforts with in-line gas analytics will be needed to elucidate this question, although were outside of the scope of this investigation.

## 5. Conclusions

*C. reinhardtii* has emerged recently as an alternative host for heterologous metabolite production using light and CO_2_ as inputs. We show here that *C. reinhardtii* can be engineered to produce significant titres of volatile isoprene in its headspace. Screening of different terpene synthases from plants directly in alga mediated by advances in synthetic transgene designs for this host allowed us to identify those which could generate robust isoprenoid yields. We show that the combination of perturbing up- and downstream pathways to DMAPP can be complementarily combined to produce significant isoprene yields from the algal cell and that temperature is likely the most important factor to improving yields beyond enzyme selection and metabolic engineering. As we relied on closed cultivation in analysis-ready headspace vials, only photo-mixotrophic growth could be analysed in this study. Future efforts will employ in-line multi-parallel analysis of culture headspaces to determine the production rates possible from this organism when grown exclusively on CO_2_. The results of our environmental parameter analysis and pathway engineering provide a strong platform for future efforts to build bio-processes with this alga to convert waste streams into value. The engineered alga has also recently been demonstrated to proliferate well in unsterile wastewaters and could be an emerging host for effluent polishing from water treatment strategies while concomitantly generating algal biomass and the volatile isoprene co-product. Algal cells producing the most volatile isoprene here also accumulate valuable ketocarotenoids as part of engineering of the downstream pathway, presenting a secondary layer of value to the biomass. These multi-product features of the cell make engineered isoprene and ketocarotenoid producing *C. reinhardtii* a very attractive system for scaled bio-refinery-processes.

## Supporting information

Plasmids sequences

Spectra of growth lights

All data from figures

## Abbreviations

IspS: Isoprene synthase
DMAPP: dimethylallyl pyrophosphate
IPP: isopentenyl-diphosphate
MEP: 2-C-methyl-D-erythritol 4-phosphate
DXS: 1-deoxy-D-xylulose 5-phosphate synthase
IDI: isopentenyl-pyrophosphate delta-isomerase

## 6. Author Contributions

RZY, GBW, and SO performed experiments and contributed to experimental design, data collection, data visualization. KJL was responsible for experimental design, project scope, funding acquisition. All authors contributed to the manuscript writing and approved the submitted manuscript version.

## 8. Acknowledgements

We would like to thank the KAUST Lab Equipment Maintenance (LEM) team and everyone involved in the setup and maintenance of the HS-GC-MS-FID, Dr. Najeh Kharbatia, Abdulkhalik M. Khalifa, and Gerard Clancy. KJL acknowledges baseline research funding provided by King Abdullah University of Science & Technology.

## 10. Supplementary Figure Legends

**Supplementary Figure 1.**
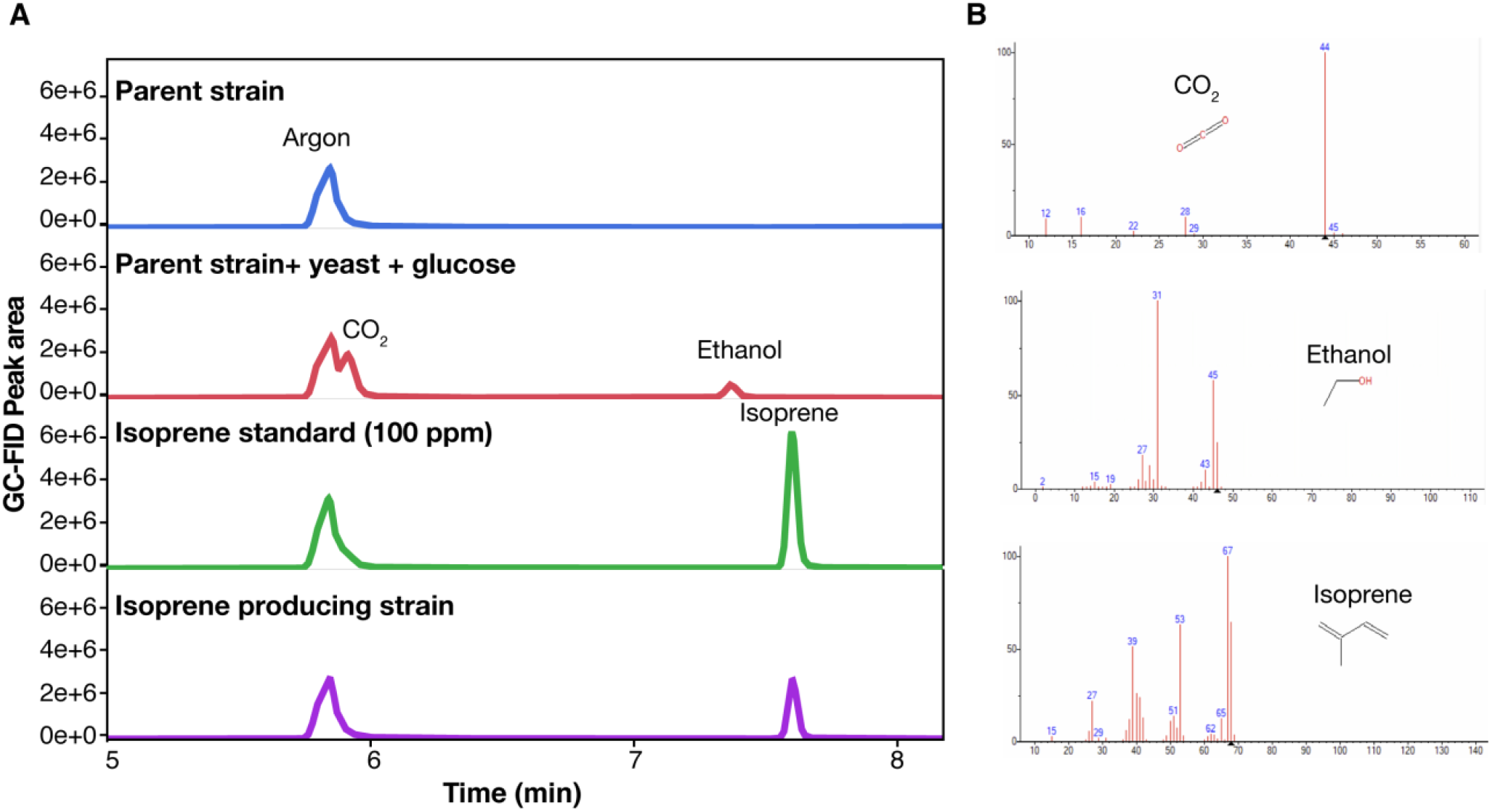
(A) GC-MS-FID chromatogram analysis of different strains showing distinct compound peaks, (B) mass spectra of compounds from peaks.

**Supplementary Figure 2.**
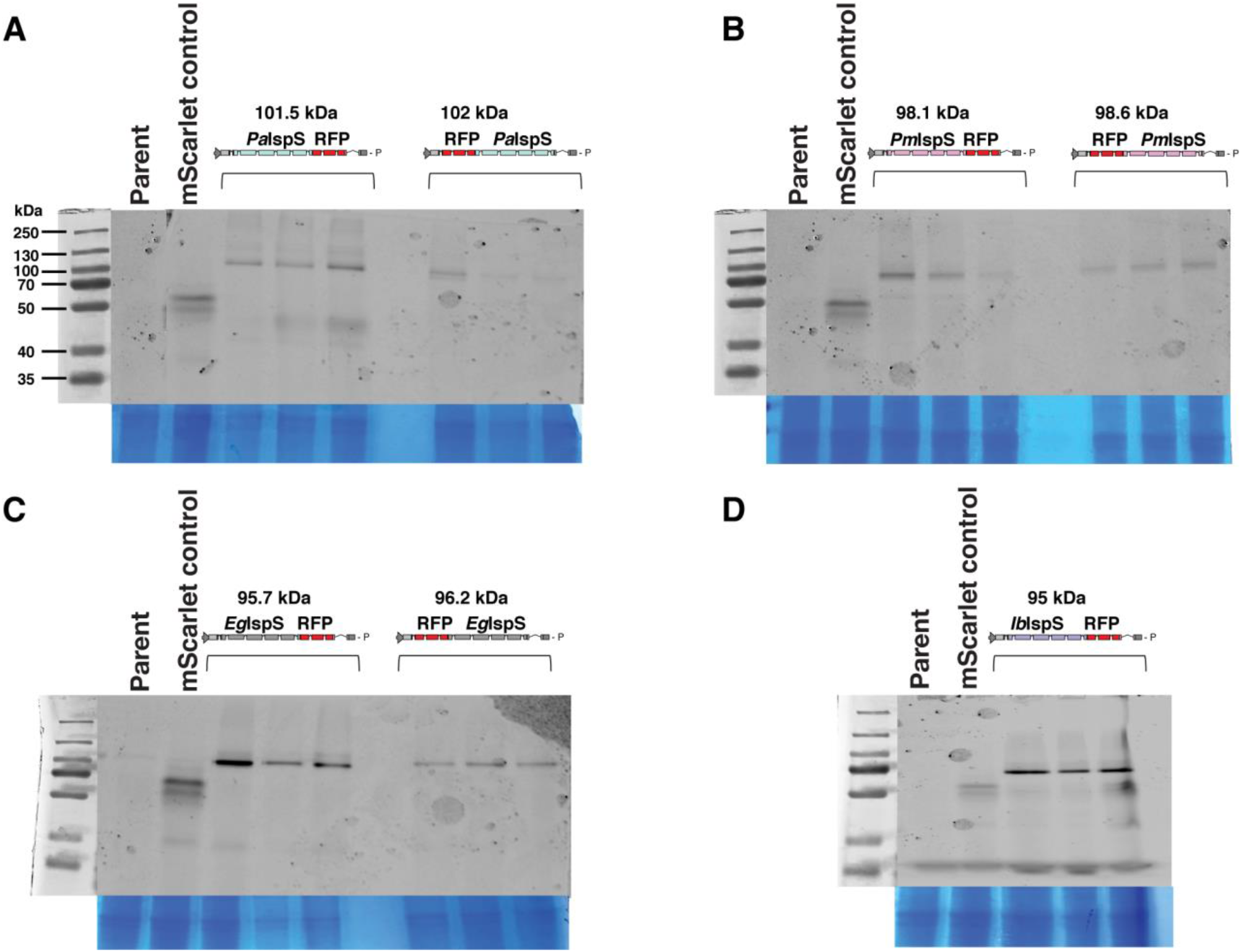
In-gel SDS-PAGE fluorescence analysis of (A) plasmid 2&6 (B) 3&7, (C) 4&8, (D) 5&9 with lanes of parental UPN22 strain (Parent) and mScarlet expressing UPN22 (RFP) control to differentiate predicted molecular masses of translated heterologous proteins.

**Supplementary Figure 3.**
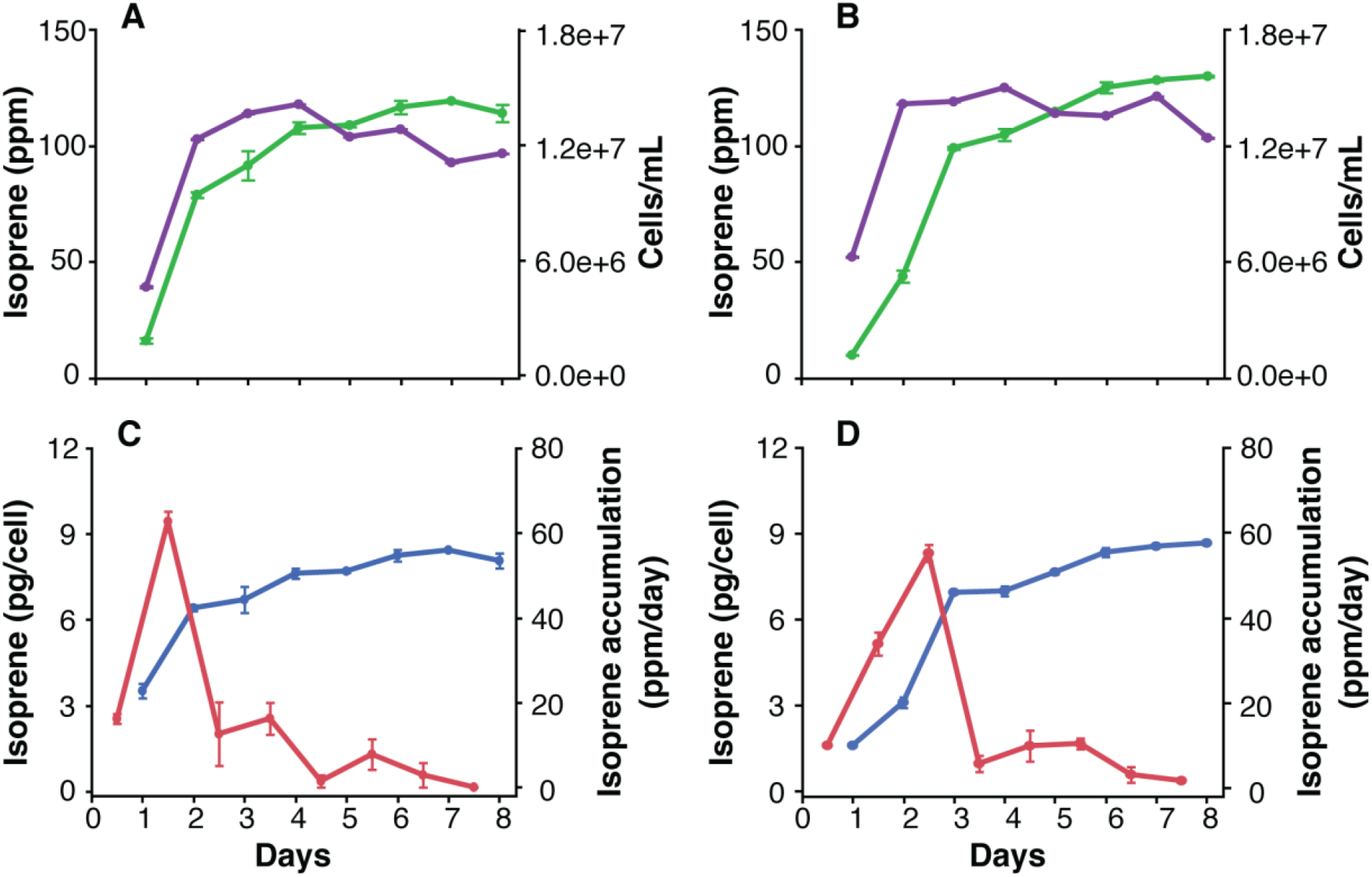
(A and B) Volumetric isoprene production and *C. reinhardtii* cell concentration, (C and D) Isoprene production (per cell) and isoprene accumulation (per day). (A and C) represent transformant 1, while (B and D) represent transformant 2 both expressing the *Ib*IspS-RFP construct from plasmid #5. Green dots represent volumetric isoprene production, purple represent cells concentration, blue dots represent isoprene production (per cell) and red dots represent isoprene accumulation (per day).

**Supplementary Figure 4.**
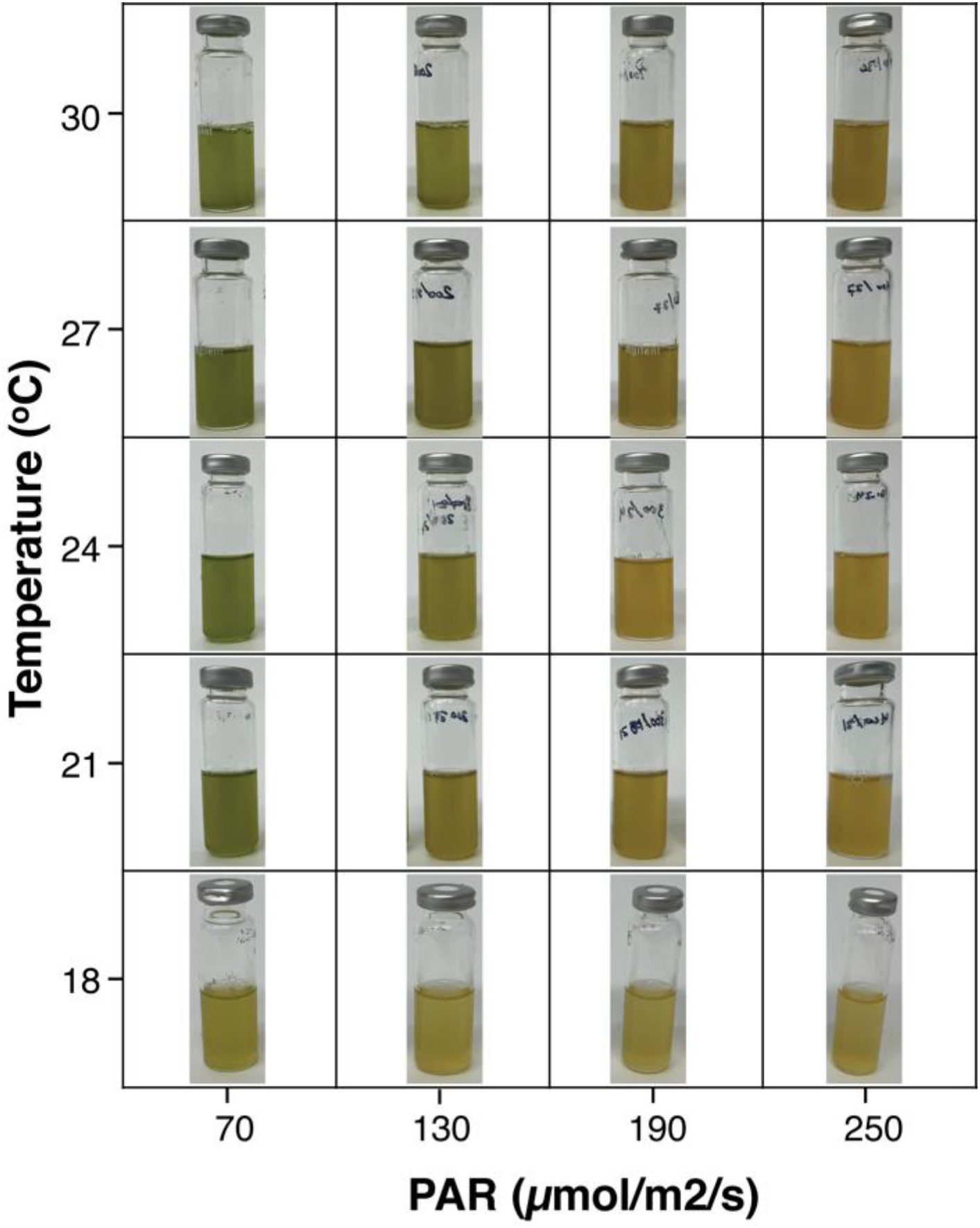
Single representative vial of cultures used to make Figure 5A. Each vial was on of 4 replicates in unique temperature and light intensity combinations achieved in controlled photobioreactors with a transformant of plasmids 5+12+25 starting from the same large pre-culture.

